# A thalamo-parietal cortex circuit is critical for place-action coordination

**DOI:** 10.1101/2022.12.30.522207

**Authors:** Christine M. Simmons, Shawn C. Moseley, Jordan D. Ogg, Madeline Johnson, Benjamin J. Clark, Aaron A. Wilber

## Abstract

The anterior and lateral thalamus (ALT) contains head direction cells that signal the directional orientation of an animal within an environment. ALT has direct and indirect connections with the parietal cortex (PC), an area hypothesized to play a role in coordinating viewer-dependent and viewer-independent spatial reference frames. This coordination between reference frames would allow an individual to translate movements toward a desired location from memory. Thus ALT-PC functional connectivity would be critical for moving toward remembered locations. This hypothesis was tested with a place-action task that requires associating an appropriate action (left or right turn) with a spatial location. There are four arms, each offset by 90 degrees, positioned around a central starting point. A trial begins in the central starting point. After exiting a pseudorandomly selected arm, the rat had to displace the correct object covering one of two (left versus right) feeding stations to receive a reward. For a pair of arms facing opposite directions, the reward was located on the left, and for the other pair, the reward was located on the right. Thus, each reward location had a different combination of allocentric location and egocentric action. Removal of an object was scored as correct or incorrect. Trials in which the rat did not displace any objects were scored as ‘no response’ trials. After an object was removed, the rat returned to the center starting position and the maze was reset for the next trial. To investigate the role of the ALT-PC network, muscimol inactivation infusions targeted bilateral PC, bilateral ALT, or the ALT-PC network. Muscimol sessions were counterbalanced and compared to saline sessions within the same animal. All inactivations resulted in decreased accuracy. Only bilateral PC inactivations resulted in increased no response trials, increased errors, and longer latency responses on the remaining trials. Thus, the ALT-PC network is critical for linking an action with a spatial location for successful navigation.

## Introduction

It is an adaptive trait for any animal to be able to navigate through the world with purposeful goals (Gallistel, 1990; O’Keefe & Nadel, 1978). Accurate navigation is necessary to guide behavior toward places for food, away from aversive locations, toward a home base from a recently learned location, or when exploring new places that were recently discovered. Goal locations are always changing depending on the needs of an animal and requires brain circuitry that can support these navigational needs in a flexible and adaptable manner. A large body of research has shown that a cortical-limbic circuit is responsible for representing the spatial layout of the environment in a map-like (allocentric) frame of reference, thereby supporting accurate spatial navigation (McNaughton, Battaglia, Jensen, Moser, & Moser, 2006; O’Keefe & Nadel, 1978; O’Mara & Aggleton, 2019; Zhao, 2018). However, interacting with the environment on one level are centered on the position of the animal’s body in that environment or fixed to the animals view of the environment; therefore, the brain must represent space with respect to spatial frames of reference that can support different perspectives of the environment (Wang, 2012). Research has provided evidence that a variety of brain regions support the use of different spatial reference frames (Alexander et al., 2022; Clark, Simmons, Berkowitz, & Wilber, 2018; Nitz, 2006, 2009, 2012; Ormond & O’Keefe, 2022; Wilber, Clark, Forster, Tatsuno, & McNaughton, 2014). Two well-studied reference frames are known as egocentric and allocentric coordinate systems. Where a goal location respective to the self is considered “egocentric” and goal location relative to landmarks is considered “allocentric” (Byrne & Crawford, 2010). Studies have employed different tasks (y-maze, Morris water maze, and the radial arm maze) to isolate and identify neural circuits and behaviors that underlie egocentric and allocentric reference frame use in navigation. However, egocentric and allocentric perspectives or strategies do not always work independently from one another, but can work in tandem and, therefore, it can be difficult to isolate their respective contributions on spatial behavior (McDonald & White, 1994; Sutherland & Hamilton, 2004; Whishaw, Hines, & Wallace, 2001). Similarly, the brain circuits that underlie the use of different strategies overlap in their function (e.g., the parietal cortex and retrosplenial cortex contain cells that encode in allocentric or egocentric reference frames or both; see Alexander et al., 2022; Nitz, 2006, 2009, 2012; Wilber et al., 2014).

The rodent parietal cortex (PC) has been shown to contain both single cells and modules (large groups of adjacent cells with consistent encoding across depth) encoding of motion states such as running straight forward at a particular speed or turning at a particular angular velocity and encoding of 3D body position (Mimica, Dunn, Tombaz, Bojja, & Whitlock, 2018; Nitz, 2006, 2009, 2012; Whitlock, Pfuhl, Dagslott, Moser, & Moser, 2012; Wilber et al., 2014; Wilber, Skelin, Wu, & McNaughton, 2017). However, the PC region has a heterogenous representation of space, in that it has cells that respond to egocentric or allocentric representations or both (Nitz, 2009; Wilber et al., 2014). For instance, in a task where rat are trained to run to randomly ordered set of cue lights, recorded cells in the PC were found to be modulated by egocentric cue direction, allocentric head direction, or a conjunctive combination of this information (Wilber et al., 2014). In addition, when the PC is damaged, animals exhibit severe navigation deficits such that the the path taken to goal locations is usually inefficient (reviewed in: Clark et al., 2018; Kolb & Walkey, 1987). Thus, the PC has a role in guiding accurate navigation toward goal locations.

The PC receives extensive input from the anterior and lateral thalamic nuclei; both of which are thought to have a critical role in processing spatial information for navigation (Aggleton & Nelson, 2015; Clark & Harvey, 2016; Peckford et al., 2014; Perry & Mitchell, 2019). Head direction (HD) cells, which fire as a function of an animals HD and are anchored to a fixed position in the room or environment but are also modulated by an animals self-motion (vestibular cues), are found in both the anterior and lateral thalamus, particularly the anterodorsal, anteroventral, anteromedial, and laterodorsal subnuclei (Butler, Smith, van der Meer, & Taube, 2017; Clark & Harvey, 2016; Clark & Taube, 2012; Dudchenko, Wood, & Smith, 2019; Jankowski et al., 2015; Marchette, Vass, Ryan, & Epstein, 2014; Taube, 2007; Yoder & Taube, 2014). The HD cell signal is also found in the retrosplenial cortex (RSC) and in other limbic-cortical areas (e.g., medial entorhinal cortex, parasubiculum, and postsubiculum). Research comparing the anchoring characteristics of HD cells suggest that distal cues are likely to modulate their activity more so than proximal/foreground cues (Knight & Hayman, 2014). This is likely due to the relative permanence of background cues. Further, studies have shown that damage to the anterior-lateral thalamic nuclei (ALT) impairs the acquisition and retention of allocentric information, but does not impair navigation based on egocentric information or visual cues (Aggleton & Nelson, 2015; Clark & Harvey, 2016; Harvey, Thompson, Sanchez, Yoder, & Clark, 2017; Lopez et al., 2009; Moreau et al., 2013; O’Mara, 2013; Peckford et al., 2014; Wolff, Gibb, Cassel, & Dalrymple-Alford, 2008). These findings demonstrate a critical role for the ATL region in the navigational ability that relies on an allocentric strategy.

In many computational and theoretical models, the thalamic HD signal is critical for translating between allocentric and egocentric coordinate systems, as an animal’s heading position is required to know the relationship between the self and the world (Bicanski & Burgess, 2016, 2018; Byrne, Becker, & Burgess, 2007; Calton, Turner, Cyrenne, Lee, & Taube, 2008; Clark, Bassett, Wang, & Taube, 2010; Clark & Taube, 2012; Pouget, Deneve, & Duhamel, 2002). The ALT and PC are anatomically and functionally connected and are in a prime anatomical position to serve as a translational interface between egocentric and allocentric frames of reference (Wilber et al., 2015). Although both structures contain HD cells (Taube, 1995; Wilber et al., 2014), a fundamental coding scheme in the PC is action centered (Wilber et al., 2017), positioning it as a critical structure for interfacing between allocentric representations and action. Anatomically, connections exist between ALT and the PC both directly and indirectly via the RSC (Clark et al., 2018; Wilber et al., 2015). Thus, the ALT-PC or ALT-RSC-PC circuit may be critical for interfacing between action centered and allocentric frames of reference. The present study was aimed at testing this anatomical and theoretical hypothesis using a novel place-action task (similar to: Grieves, Jenkins, Harland, Wood, & Dudchenko, 2016) and muscimol induced disconnection of the ALT and PC. Briefly, the place-action task requires that rats perform a specific action when at a specific orientation/place in the environment. Thus, we specifically hypothesize that disruption of functional connectivity between dorsal-medial thalamus and PC will impair performance in this task. Functional disconnection was performed by selectively inactivating the thalamus and PC contralaterally using muscimol infusions targeting each region in one hemisphere (e.g. right PC and left ALT; Fresno, Parkes, Faugère, Coutureau, & Wolff, 2019; Jo & Lee, 2010). Ipsilateral infusions are used as a control because the pathway in the opposite hemisphere is left intact. For the current work, the functional connectivity between the ALT and the PC was disrupted using the same inactivation technique. The ALT has dense ipsilateral (but not contralateral) projections to PC, which does not have many reciprocal connections to ALT (Wilber et al., 2015). Thus, we took advantage of the primarily ipsilateral anatomical connectivity between ALT and PC to disrupt this circuit with contralateral or “cross” infusions. We designed a task that requires an animal to execute a specific egocentric action at each of four allocentric locations so that we could assess the function of the PC, ALT, and ALT-PC network in egocentric and allocentric coordination.

## Methods

Subjects were 6 female and 5 male Long Evans rats between 2 and 11 months old. All animals were housed individually throughout the experimentation process in a 12h lights on light and dark cycle. For all behavioral training and experimentation, rats were food deprived to no less than 80% of their ad libitum body weight. Rats were given full access to water for all phases of experimentation. All procedures carried out were in accordance with the NIH Guide for the Care and Use of Laboratory Animals and approved by the Florida State University Animal Care and Use Committee.

### Pre-training

The maze apparatus was secured on top of a circular arena measuring 5 feet in diameter and consisted of 4 walled pathways or “arm” leading out from a central chamber, with 1 door leading to each arm, and each arm positioned at a 90-degree angle from the adjacent arms (**Fig. 1A**). Each pathway was parallel to a pair of walls in the room. This layout created 4 paths, or arms, of equal length that were centered around the middle of the circular arena. The width of the arms was equivalent to the width of each door such that when all doors were closed, a square region in the center of the arena was partitioned off from the 4 arm entry points; this center region was the starting point for each trial. At the end of each arm, a weigh boat was secured to the arena’s surface and covered with a disc. For pretraining rats were progressively trained to run through a maze arm and remove a plastic disc that rested on top of a small, square weigh boat.

**Figure 1.**
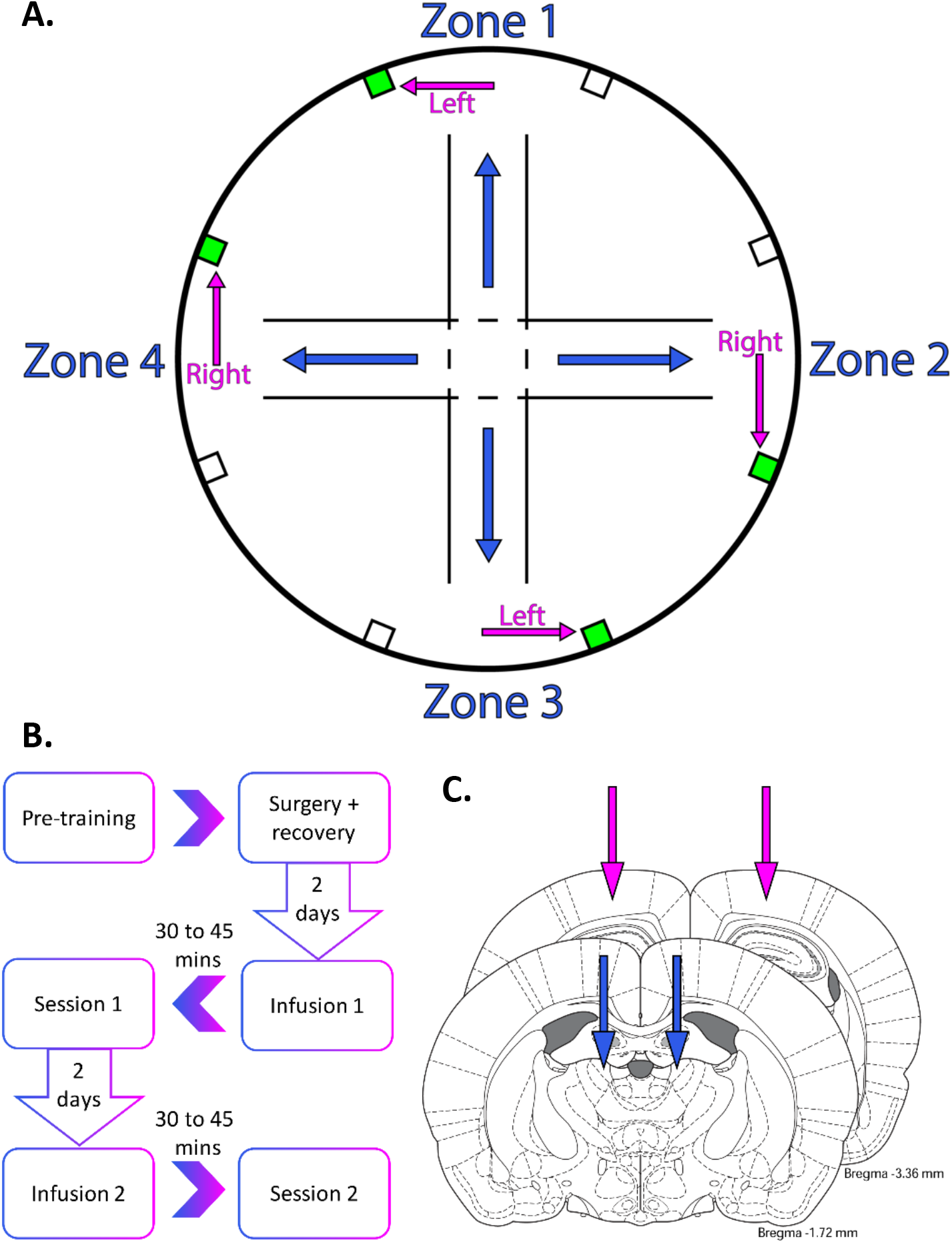
Place-action apparatus, experimental timeline and surgical targets. **A.** Four arms of equal length each bisect a pair of feeding stations - one rewarded, ‘correct’ and one non-rewarded, ‘incorrect’ location. Each reward station is associated with a different combination of allocentric place and egocentric action. **B.** Behavioral sessions and pharmacological inactivation timeline. Projections between the ALT and PC are laregly ipsilateral, and so the network can be tested with contralateral inactivations. **C.** Double bilateral cannulae targets: PC (magenta) and ALT (blue).

For the 1st phase of pre-training, the doors remained open and 2 frootloops were placed in each weigh boat under the discs. The rats began each session at the center starting point and were allowed to explore the maze with little intervention. Whenever the rats removed a disc to eat a frootloop underneath, the experimenter re-covered the weigh boat containing the remaining frootloop. Once a weigh boat was emptied, it was refilled with 2 more frootloops. Each 1^st^ phase pre-training session was conducted for 20 minutes every day until the rat removed the disc 5 times from each weigh boat.

In the second phase of pre-training, the doors remained open, but the weigh boats were never filled with frootloops. Rats were hand fed a frootloop after removing any disc from any weigh boat. Each 2^nd^ phase pre-training session was conducted for 20 minutes and continued every day until the rats removed discs 5 times from each weigh boat.

In the 3rd and final phase of pre-training, the sessions were conducted as they were in the 2nd phase pre-training, except that only 1 door, selected randomly by the experimenter, was open to a single arm per trial. The rats were trained to run down an open arm, remove the disc at the end of the arm, and return to the center starting point after receiving a reward at the location of the removed disk. This was repeated for the entire 20-minute session. The 3^rd^ phase pre-training sessions continued every day until the rat again removed discs 5 times from each weigh boat.

### Place-Action Task

The maze apparatus was set up as described above. The full task was conducted as described in the 3^rd^ pretraining phase, but now 8 disc-covered weigh boats that were offset to the left and right sides of each arm and secured closer to the edge of the arena (**Fig. 1A**). Each pair of weigh boats, bisected by an arm, consisted of a ‘correct’ reward zone and an ‘incorrect’ zone. For two of these weigh boat pairs, the reward zone was located on the right side of the arm and for the other two pairs, the reward zone was located on the left side of the arm. Each of the reward zones were assigned a numerical label (1, 2, 3, 4) and were input into a random list generator (Random.org) in order to generate a randomized 40-item list containing each reward zone number exactly 10 times. The list was then rearranged manually to avoid single (4-4) and double (1-2-1-2) zone repeats. This pseudorandomized list was generated for each behavioral session and determined the arm order for each of the 40 trials.

The rats began every trial in the center of the maze. To start each trial, a door was opened leading into the pseudo-randomly selected arm. Rats were rewarded for removing the disc covering the reward well for the correct side. If a rat removed a disc from the incorrect side or did not remove a covering after 1 minute, these trials were marked as incorrect and no response trials (NR) respectively. After a correct (C) response, incorrect response (I), or NR, the trial was completed, and returned to the center starting point. The experimenter would record the response on the trial list, clean the traversed area with ethanol, exchange the rewarded and incorrect zone covers, and begin the next trial. The session ended when all 40 trials on the list were completed. The rats were scheduled for cannula implantation surgery once they reached a criterion performance of at least 85% correct for two consecutive days. The animals were given full access to food after this criterion was met and scheduled for surgery.

### Cannula Implantation Surgery

All 11 rats were surgically implanted with two sets of cannula bilaterally targeting both the PC (anterior-posterior −4.5 mm, medial-lateral ± 3 mm, dorsal-ventral −0.1 mm) and the ALT (anterior-posterior −1.74 mm, medial-lateral ± 1.25 mm, dorsal-ventral −5.23 mm). After surgery, all animals were given 7 days to recover with full access to food and water.

### Infusion and Behavior Timeline

After the post-surgical recovery period, rats continued the behavioral sessions as previously described in the Place-Action Task methods. When the animal again reached the criterion performance of at least 85% correct for two consecutive days, the animal was scheduled to receive the first infusion the next day. For the first infusion, saline or muscimol (order was decided randomly and pairs were counterbalanced) was infused bilaterally into the PC before the behavioral session (**Fig. 1B-C**). For the days following the infusion day, rats continued the behavioral task until criterion performance was again reached. The animal was then scheduled to receive the second infusion the next day. The second infusion was repeated with the counterbalanced solution (saline or muscimol) before the behavioral session. This entire procedure is repeated until the bilateral PC, contralateral PC and ALT pairs (Left PC + Right ALT, Right PC + Left ALT), and bilateral ALT all received successful pairs of muscimol and saline infusions, unless one of the infusion sites became obstructed first (for example by dura regrowth). All ALT infusions occurred 30 minutes before a behavioral session whereas all PC infusions occurred 45 minutes before a behavioral session. This timing difference was based on the differential sizes of the two structures and pilot/experimental data suggesting these timings were optimal for ensuring complete spread that was not likely to encroach on adjacent structures. The behavioral performance between the saline and muscimol infusion days were then compared for each pair of sessions.

### Infusion Details

All infusions were done with two 10-ml, 22-gauge Hamilton syringes held in a Model ‘22’ Harvard Apparatus syringe pump. The rats were lightly restrained, and the dummy cannula were removed before inserting the infusion cannula into the infusion guides (dual bilateral – PC and ALT). All PC infusions were done at a rate of 0.3 ml/min for 1 min 45 min before the behavior session began. Infusion cannula were kept inside the guides for 1 min after the infusion before removing and reinserting the dummy cannula. All ALT infusions were done at a rate of 0.167 ml/min for 1.5 mins 30 min before the behavior. ALT infusion cannula were kept inside the guides for 30 s after the infusion and then removed before reinserting the dummy cannula. For the network infusions, a unilateral PC infusion was completed 15 minutes prior to a subsequent unilateral ALT infusion, which was always contralateral to the hemisphere that received the PC infusion. Again, differences between ALT and PC infusion parameters were due to the smaller structure to optimize sufficient time for coverage of the structure but prior to significant diffusion into adjacent structures.

### Statistics

Separate repeated measures (RM) ANOVAs were performed for numbers of trials and percentages. Group main effects (muscimol vs saline) are not reported for any of the infusion targets (PC, ALT, ALT-PC) because there was no significant difference in numbers of trials within a session. RM ANOVAs were followed by planned comparisons. Planned comparisons consisted of two-group *t*-tests done within the context of the overall ANOVA (Maxwell & Delaney, 2003), comparing the saline condition to the inactivation condition for each performance category (C, I, NR). For all statistical analyses, p < 0.05 was considered significant and the soft-ware used for statistical analyses was StatView (SAS Institute Inc.). Paired t-tests within each region were used to assess for possible effects of muscimol on trial duration, side bias, head scanning, and procedural errors.

### Histology

Once all sets of infusions (PC, ALT, ALT-PC and unilateral) and behavioral sessions were completed, the rats infused with fluorescent muscimole or labeled AAV into PC and ATN and then were deeply anesthetized with an IP injection of a sodium pentobarbital solution, then perfused transcardially with a 1X phosphate-buffered saline solution (PBS) followed by a 4% paraformaldehyde (PFA) in 1X PBS solution. The whole head was removed and preserved in the 4% PFA solution for 24 h before extracting the brain and fixing in 4% PFA for another 24 h. Then the brain was moved to a 30% solution of sucrose for cryoprotection. The brain was frozen and cut coronally at 40 μm thick with a sliding microtome. Slices were mounted onto slides with a mounting media containing DAPI and then the slides were coverslipped. Cannula placements were verified using a scanning microscope (Zeiss Axioimager M2).

## Results

Histological analysis revealed that most of the cannula were placed within the ALT and PC. The center point and the spread of fluorescent muscimole or AAV were recorded (**Fig. 2**). When canula were clogged before perfusion the spread was estimated by averaging the spread from all remaining animals. ATL placements typically included a combination of anterior (anterodorsal, anteroventral) and lateral thalamic (laterodorsal, centrolateral, lateral mediodorsal) nuclei. However, for 1 rat, ALT cannula were placed in the fimbria of the hippocampus. Therefore, this rat was excluded from further ALT and network analyses leaving 7 rats and 8 paired data sets for ALT only inactivation. In addition, 4 animals had PC guide cannula blockage prior to the first PC infusion, which resulted in the inability of muscimol (and saline) to diffuse into the cortical tissue. Therefore, these animals were excluded from further PC and network analyses. Thus, 7 rats and 8 paired data sets remained for PC only inactivation. Finally, due to the missed placement for ALT and additional clogging of PC canula that occurred after bilateral PC inactivation, 4 rats and 7 paired data sets remained for ALT-PC network inactivation. As a probe test for the contralateral network inactivations, unilateral inactivations were also performed to confirm that the contralateral ALT-PC inactivation behavioral changes were not the result of general inactivation of two unilateral regions, but rather, were the result of disrupting the network formed between the two regions. However, since this manipulation was performed last further increasing the likelihood that PC cannula were obstructed, only 2 paired data sets from 1 animal were included for further unilateral infusion analyses. Overall, each animal provided 1-4 datasets per inactivation condition.

**Figure 2.**
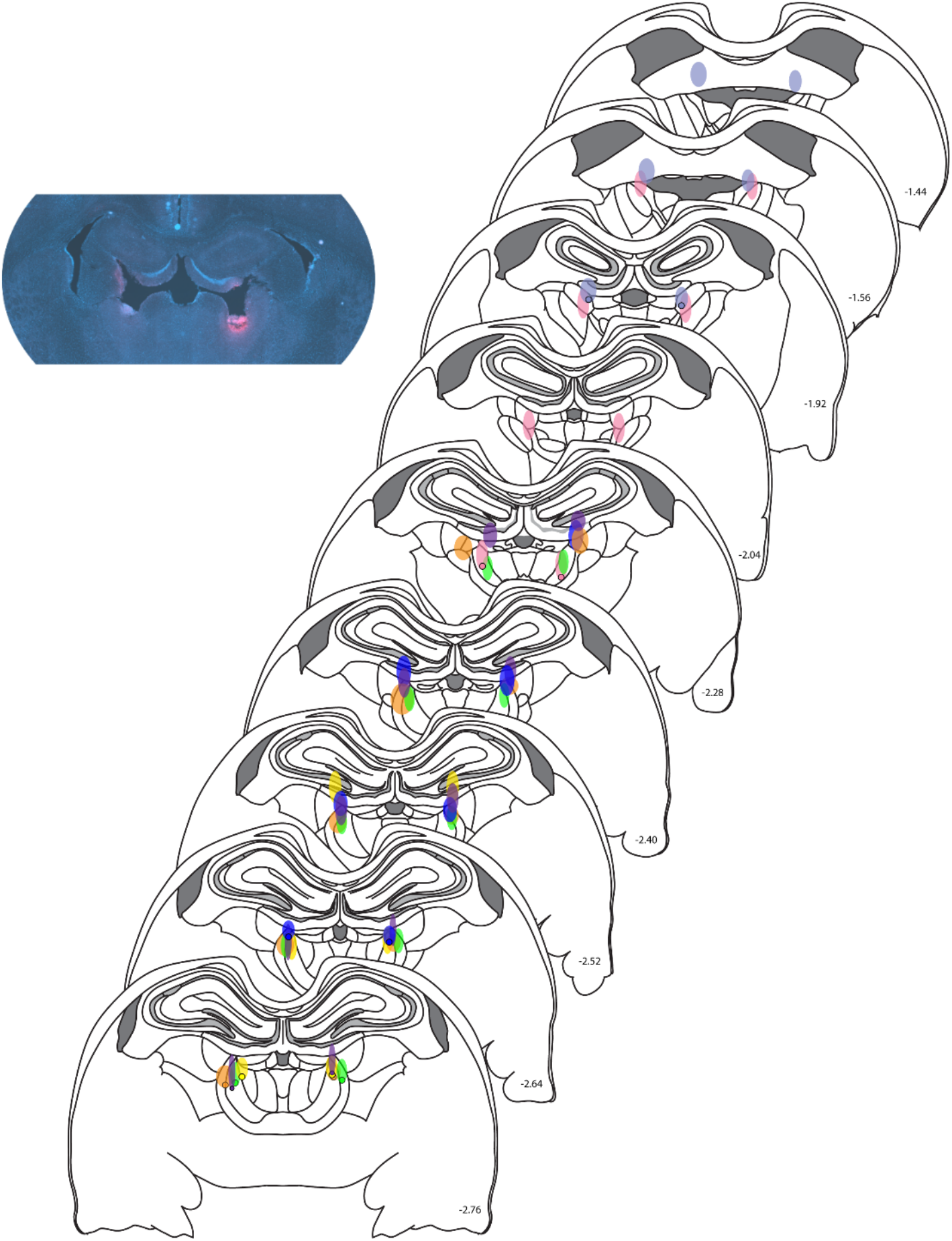
Histological verification of cannulae placements. Depiction of ALT (Middle) cannulae placement ranges overlayed on atlas images. Each color represents the verified range of fluorescent muscimol or damage of one animal. *Top Left*. Example of tissue imaged (*Left*) for verification of fluorescent muscimol presence (red signal) in ALT.

### Parietal Cortex

A RM ANOVA was performed to investigate the relationship between PC inactivation (muscimol vs saline) and performance on the place-action task (C, I, NR). Separate RM ANOVAs were performed for numbers of trials and percentages. The number of trials (not shown; F_(2,14)_ = 691.98, p < 0.0001) and percentage (**Fig. 3** *Top Left;* F_(2,28)_ = 693.27, p < 0.0001) both varied across *performance category* (main effect of performance category). Consistent with our hypothesis, *inactivation* produced variation in performance *across category* for both numbers of trials (F_(2,28)_ = 18.30, p < .0001) and percentages (F_(2,28)_ = 18.02, p < 0.0001). Specifically, PC inactivation reduced the number of correct trials (t_(7)_ = −4.46, p < 0.01), and increased both incorrect (t_(7)_ = 3.17, p < 0.05) and no response (t_(7)_ = 2.81, p < 0.05) trial counts. Similarly, PC inactivation significantly reduced the percentage of correct trials (t_(7)_ = −4.37, p<0.01), and increased both incorrect (t_(7)_ = 3.17, p < 0.05) and no response (t_(7)_ = 2.81, p < 0.05) percentage as compared to saline. In addition, the average trial duration for PC inactivation sessions was significantly longer than saline sessions (**Fig. 3** *Top Right;* t_(7)_ = 2.36, p = 0.05). Thus, PC inactivation impaired performance on the place-action task by increasing both errors and non-responding.

**Figure 3.**
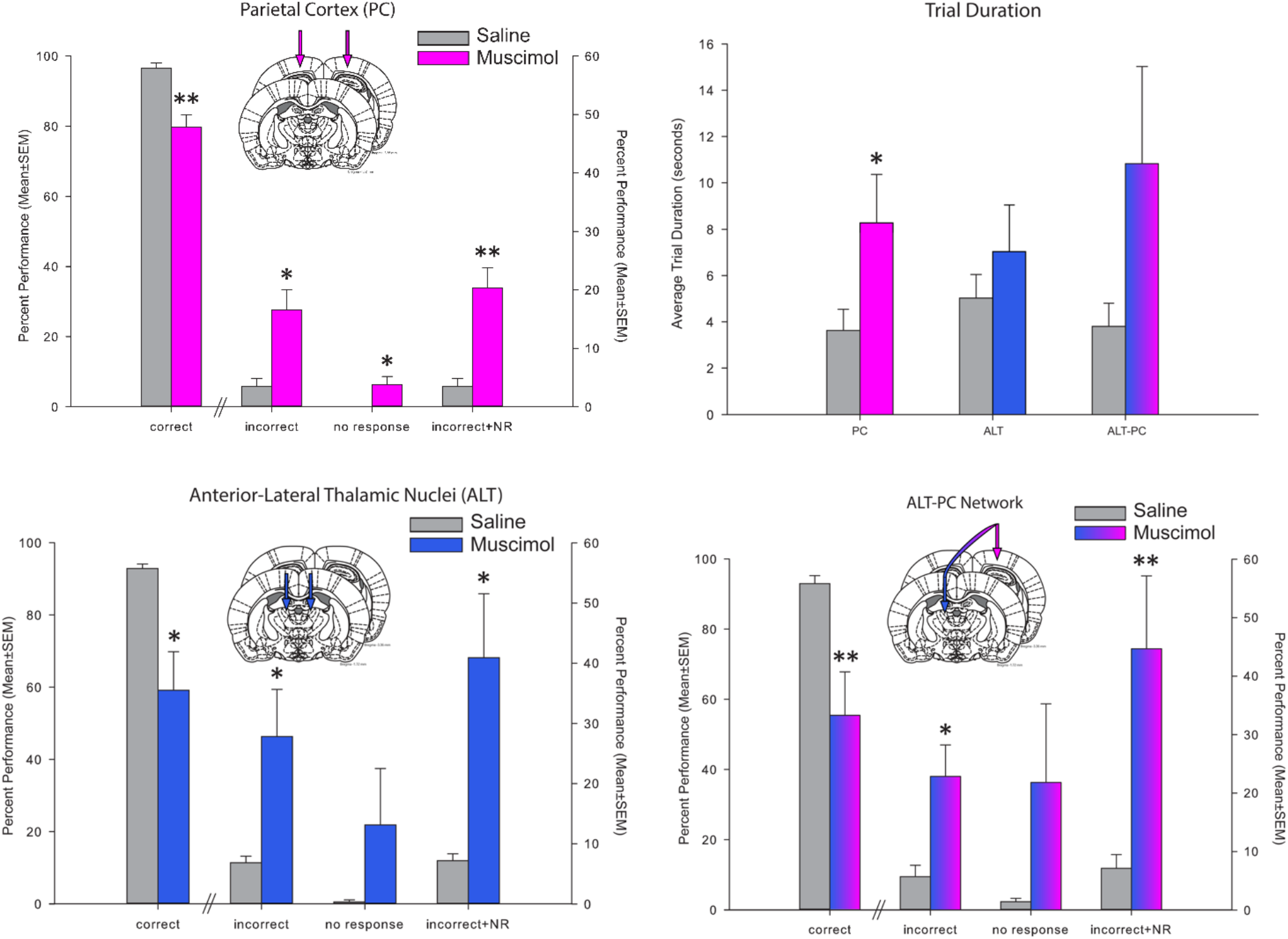
Inactivation Effects on Place-Action Task Performance. Inactivation of the PC (*Top Left*), ALT (*Bottom Left*), and the ALT-PC network (*Bottom Right*) all produce similar impairments in the task resulting in significant interactions between accuracy and performance category (Fs(2,14)> 16.7, ps < 0.05). Specifically, for each inactivation condition there was a significant reduction in percent correct and a significant increase in percent incorrect. ALT inactivation animals tend to navigate quickly to the incorrect location, potentially as a consequence of a perceived orientation that is incorrect. In contrast, PC inactivation increases the duration of each trial (*Top Right*), consistent with impaired linking of the correct action. Data set is the n (n=8 PC, n=8 ALT, & n=7 network data sets). * p < 0.05, ** p < 0.01.

To ensure that a small number of data sets were not driving the observed effects we also performed the analyses using animals as the n instead of datasets. The analyses performed above were repeated by averaging the multiple datasets for one animal when there was more than one for an inactivation condition, so that each animal had 1 dataset per inactivation condition. The number of trials (not shown; F_(2,24)_ = 686.30, p < 0.0001) and percentage (**Fig. 4** *Top* F_(2,24)_ = 688.70, p < .0001) both varied across *performance category*. Further, consistent with our hypothesis, *inactivation produced variation in performance across category* for both number of trials (F_(2,24)_ = 14.19, p < .0001) and percentages (F_(2,24)_ = 13.91, p < 0.0001). Specifically, PC inactivation reduced the number of correct trials (t_(6)_ = −3.95, p < 0.01), and increased both incorrect (t_(6)_ = 2.62, p < 0.05) and NR (t_(6)_ = 2.87, p < 0.05) trial counts. Similarly, PC inactivation significantly reduced the percentage of correct trials (t_(6)_ = −3.88, p<0.01), and increased both incorrect (t_(6)_ = 2.62, p < 0.05) and no response (t_(6)_ = 2.87, p < 0.05) percentage as compared to saline. In addition, the average trial duration for PC inactivation sessions was significantly longer than saline sessions (not shown; t_(6)_ = 3.16, p < 0.05). Thus, PC inactivation impaired performance on the place-action task by increasing both errors and non-responding.

**Figure 4.**
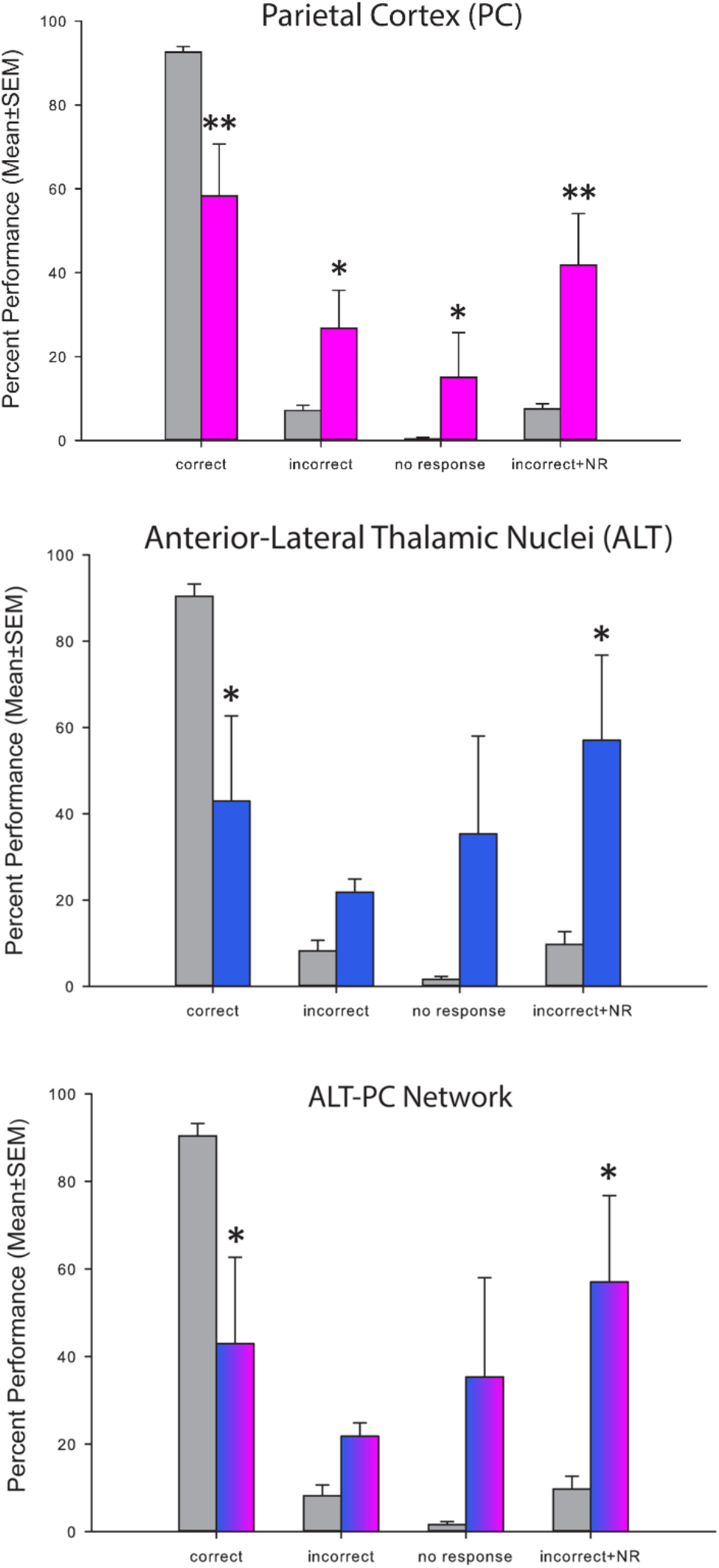
Place-action task performance with animal as n. Inactivation of the PC (*Top*), ALT (*Middle*), and the ALT-PC network (*Bottom*) all produce similar impairments to those observed with data sets as the n (n=7 animals for PC, n=7 animals for ALT, & n=4 animals for network). Specifically, for each inactivation condition there was a significant reduction in percent correct and a significant increase in the combined percent incorrect + no response. * p < 0.05, ** p < 0.01.

### Anterior-Lateral Thalamus

RM ANOVAs were also performed for numbers of trials and percentages to investigate the relationship between ALT inactivation and place-action task performance. The number of trials (not shown; F_(2,26)_ = 41.88, p < 0.0001) and percentage (**Fig. 3** *Bottom Left;* F_(2,26)_ = 41.91, p < .0001) both varied across *performance category*. Consistent with our hypothesis, *inactivation* produced variation in performance *across category* for both numbers of trials (F_(2,26)_ = 6.59, p < 0.01) and percentages (F_(2,26)_ = 6.64, p < 0.01). Specifically, ALT inactivation reduced the number of correct trials (t_(7)_ = −3.34, p < 0.05) and increased incorrect (t_(7)_ = 2.71, p < 0.05) trial counts. Unlike PC, inactivation of ALT did not significantly alter performance on no response trial counts (t_(7)_ = 1.45, p = 0.19). Results for percentages were identical to trial counts, ALT inactivation significantly reduced the percentage of correct trials (t_(7)_ = −3.43, p=0.01) and increased incorrect percentage as compared to saline (t_(7)_ = 2.72, p < 0.05) but not no response percentage as compared to saline (t_(7)_ = 1.45, p = 0.19). Unlike PC, the average trial duration for ALT inactivation sessions was not significantly different than saline sessions (**Fig. 3** *Top Right;* t_(6)_ = −1.04, p = 0.34). Thus, ALT inactivation impaired performance on the place-action task by increasing incorrect responses but not non-responding.

To ensure that a small number of data sets were not driving the observed effects we also performed the ALT analyses animals as the n. The number of trials (not shown; F_(2,24)_ = 35.64, p < 0.0001) and percentage (**Fig. 4** *Middle;* F_(2,24)_ = 35.67, p < .0001) both varied across *performance category*. Again, *inactivation* produced variation in performance *across category* for both numbers of trials (F_(2,24)_ = 5.81, p < .01) and percentages (F_(2,24)_ = 5.87, p < 0.01). Specifically, ALT inactivation reduced the number of correct trials (t_(6)_ = −2.94, p < 0.05). Unlike PC, inactivation of ALT did not significantly alter the number of incorrect trials (t_(6)_ = 2.24, p = 0.07) or no response trial counts (t_(6)_ = 1.47, p = 0.19). Results for percentages were identical to trial counts, ALT inactivation significantly reduced the percentage of correct trials (t_(6)_ = −3.03, p<0.05), but did not affect incorrect percentage (t_(6)_ = 2.24, p = 0.07) or no response percentage as compared to saline (t_(6)_ = 1.46, p = 0.19). Unlike PC, the average trial duration for ALT inactivation sessions was not significantly different than saline sessions (not shown; t_(5)_ = −0.86, p = 0.43). Thus, ALT inactivation impaired performance on the place-action task by reducing correct responses but not altering the incorrect or no response behaviors.

### Parietal-Anterior-Lateral Thamalic Network

RM ANOVAs were also performed for numbers of trials and percentages to investigate the relationship between contralateral ALT-PC inactivations and place-action task performance. The number of trials (not shown; F_(2,24)_ = 31.15, p < 0.0001) and percentage (**Fig. 3** *Bottom Right;* F_(2,24)_ = 31.15, p < .0001) both varied across *performance category*. Further, consistent with our hypothesis, *inactivation* produced variation in *across performance category* for both numbers of trials (F_(2,24)_ = 6.59, p < 0.01) and percentages (F_(2,24)_ = 6.59, p < 0.01). Specifically, ALT inactivation reduced the number of correct trials (t_(6)_ = 3.87, p < 0.01) and increased incorrect (t_(6)_ = 3.20, p < 0.05) trial counts. Unlike PC, the ALT-PC network inactivations did not significantly alter performance on no response trial counts (t_(6)_ = 1.68, p = 0.14). Results for percentages were identical to trial counts, network inactivation significantly reduced the percentage of correct trials (t_(6)_ = −3.87, p<0.01) and increased incorrect percentage as compared to saline (t_(6)_ = 3.20, p < 0.05) but not no response percentage as compared to saline (t_(6)_ = 1.68, p = 0.14). Unlike PC, the average trial duration for ALT-PC inactivation sessions was not significantly different than saline sessions (**Fig. 3** *Top Right;* t_(5)_ = −1.93, p = 0.11). Thus, ALT-PC network inactivation impaired performance on the place-action task by increasing incorrect responses but not non-responding. In summary, ALT and network inactivation produced an identical pattern of results that differed slightly from PC inactivation in that only PC inactivation increased no response trials.

To ensure that a small number of data sets were not driving the observed effects we also performed the network analyses using animals as the n. The number of trials (not shown; F_(2,12)_ = 9.60, p < 0.01) and percentage (**Fig. 4** *Bottom;* F_(2,12)_ = 9.60, p < .01) both varied across *performance category*. Finally, consistent with our hypothesis, *inactivation* produced variation in performance *across category* for both numbers of trials (F_(2,12)_ = 5.12, p < 0.05) and percentages (F_(2,12)_ = 5.12, p < 0.05). Specifically, network inactivation reduced the number of correct trials (t_(3)_ = −3.22, p < 0.05). Unlike PC, inactivation of the network did not significantly alter incorrect trial counts (t_(3)_ = 3.01, p = 0.06) or no response trial counts (t_(3)_ = 1.77, p = 0.18). Results for percentages were identical to trial counts, ALT inactivation significantly reduced the percentage of correct trials (t_(3)_ = −3.22, p<0.05), but not incorrect percentage (t_(3)_ = 3.01, p = 0.06) or no response percentage as compared to saline (t_(3)_ = 1.77, p = 0.18). Unlike PC, the average trial duration for ALT-PC inactivation sessions was not significantly different than saline sessions (not shown; t_(2)_ = −1.22, p = 0.35). Thus, network inactivation impaired performance on the place-action task by increasing incorrect responses but not non-responding.

### Unilateral Control Conditions

We assessed performance accuracy means for the unilateral infusion datasets. With the muscimol infusions into the ipsilateral regions of ALT and PC, the mean percentage correct was 88.8%. For the saline infusions, the mean percentage correct was 93.8% suggesting that unilateral inactivation did not produce the same effect as network inactivation (61.3% and 93.8% for network, muscimol and saline respectively).

### Side bias, head scanning, and procedural errors

We also investigated whether muscimol inactivation resulted in general changes in behavioral performance by measuring the number of times the animal engaged in stereotyped behaviors relating to errors or inefficiencies in navigation and orientation. Side bias, or the ratio of preferred left or right turns toward the goal location was calculated and compared between muscimol inactivation and saline control for each performance category. The side bias was calculated by taking the absolute value of the total number of left choices minus the total number of right choices and then dividing by the total number of trials. Paired t-tests were performed to assess side bias ratios for each inactivation type (PC, ALT, and network) comparing the paired saline vs muscimol data sets. There were no significant differences in side bias between muscimol and saline for any of the brain regions with either the data set as the n (**Fig. 5** *Top Left;* PC: t_(5)_ = 1.58, p = 0.18; ALT: t_(5)_ = 1.90, p = 0.12; Network: t_(5)_ = 1.68, p = 0.15) or the animal as the n (**Fig. 5** *Bottom Left;* PC: t_(5)_ = −2.09, p = 0.09; ALT: t_(5)_ = −1.70, p = 0.15; Network: t_(2)_ = −1.13, p = 0.37). Additionally, we assessed head scanning (the number of times an animal moved its head to and from the location of the goal). Paired t-tests were also used to assess the relationship between inactivation and head scanning. There were no significant differences in head scanning between muscimol and saline for any of the brain regions with either the data set as the n (**Fig. 5** *Top Middle;* PC: t_(5)_ = 0.57, p = 0.59; ALT: t_(5)_ = −1.56, p = 0.18; Network: t_(5)_ = 1.00, p = 0.36) or the animal as the n (**Fig. 5** *Bottom Middle;* PC: t_(5)_ = −0.68, p = 0.53; ALT: t_(5)_ = 1.44, p = 0.21; Network: t_(2)_ = −1.82, p = 0.21). Lastly, we looked at procedural errors which included the number of times an animal traveled around the perimeter of the arena and past any of the three other arms. There were no significant differences in the average procedural errors between muscimol and saline for any of the brain regions using session means as n (**Fig. 5** *Top Right;* PC: t_(5)_ = −2.43, p = 0.06; ALT: t_(5)_ = −1.50, p = 0.19; Network: t_(5)_ = −1.64, p = 0.16). However, there was a significant difference in the average procedural error count using animal means as n for PC inactivations but not for ALT and ALT-PC network inactivations (**Fig. 5** *Bottom Right;* PC: t_(5)_ = −2.64, p < 0.05; ALT: t_(5)_ = −1.55, p = 0.18; Network: t_(2)_ = −1.19, p = 0.36).

**Figure 5.**
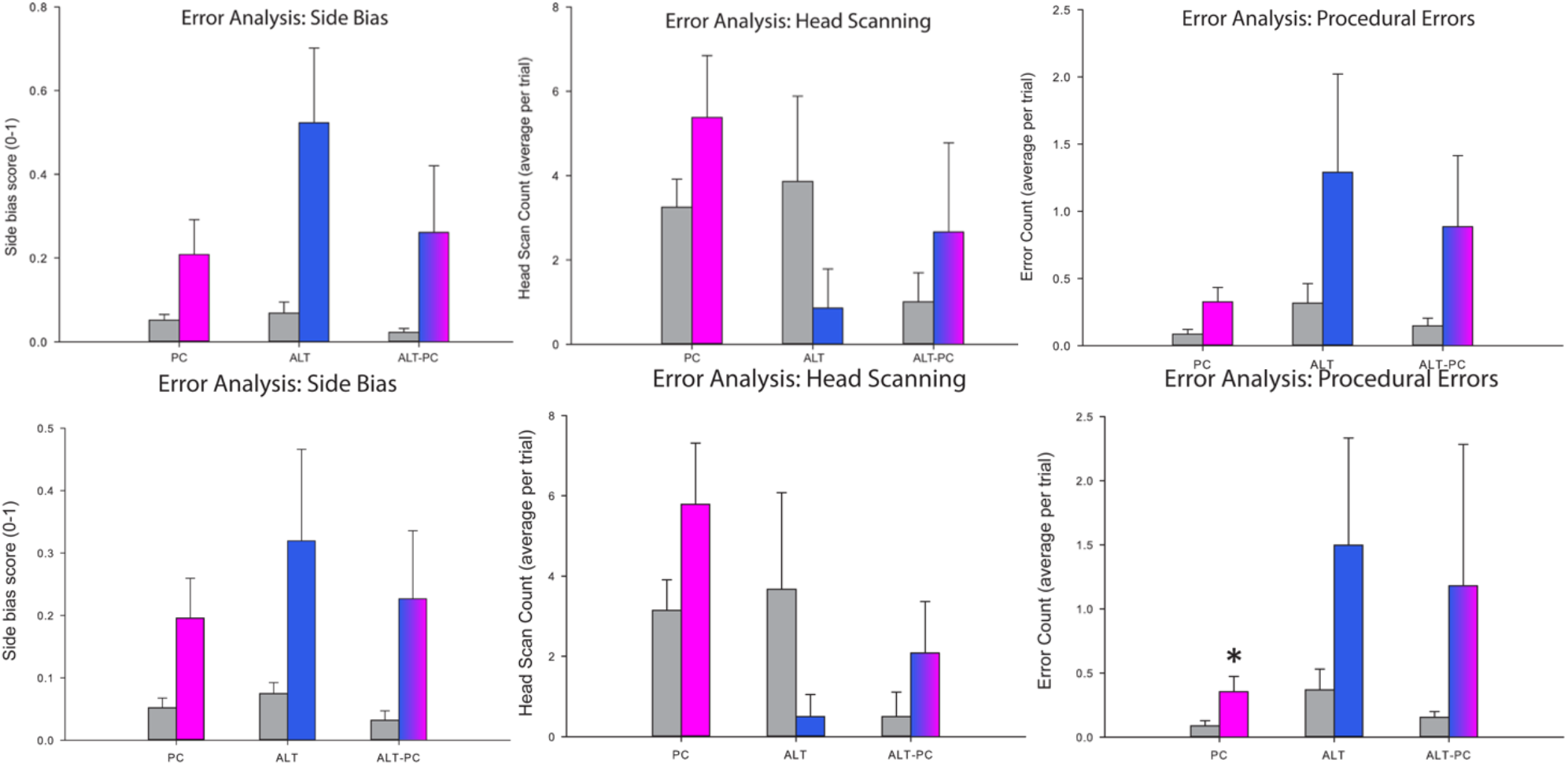
Error analysis. Error analyses are shown for session as the n (*Top Row*) and animals as the n (*Bottom Row). Left Column*. Side bias scores were calculated for each session and averaged across animals. 0 is no preference for left of right turns to target location, 1 is complete preference for one turn direction to target. Side bias did not differ for any inactivation condition (ps>0.12). *Middle Column*. The number of head scan movements made before a correct or incorrect decision averaged for each session did not differ for any inactivation condition (ps>0.19). Right Column. The mean number of procedural errors per session was significantly increased following PC inactivation but only for animals as the n. * p<0.05.

## Discussion

The purpose of the present study was to test the hypothesis that functional connectivity between the ALT and PC is necessary for linking actions with allocentric spatial information. Overall, the results demonstrated that the ability to accurately perform the place-action task decreased significantly with muscimol inactivations across all inactivation types (ALT, PC, network). Though effects were generally similar across inactivation conditions, one key difference was that only PC inactivation increased trial length, no-response trials, and procedural errors; suggesting PC is essential for generating the appropriate action to the goal location (i.e., with ALT intact but PC inhibited the rat has difficulty generating an action but with ALT inhibited PC generates the wrong action). Together this data suggests that the ALT-PC circuit is critical for transforming an allocentric location into the appropriate action.

The results of the present study are consistent with the notion that the PC has a critical role in processing egocentric and allocentric information and potentially serves as a convergence point for these two sources of spatial information. Supporting this view are results from previous studies showing that single cell encoding in the PC is mixed with both egocentric and allocentric encoding (including conjunctive encoding in both reference frames); however, mesoscale encoding (multi-unit activity) in PC is organized around motion state (Kolb, Buhrmann, McDonald, & Sutherland, 1994; Kolb & Walkey, 1987; Wilber et al., 2014; Wilber et al., 2017). Further, motion state encoding in PC is sometimes anticipatory, predicting the upcoming action (Wilber et al., 2014). This encoding scheme at the single unit scale and mesoscale combined with the present results suggests PC plays a critical role in the ability to access an allocentric map and use this information to navigate towards a desired goal. Therefore, the present finding of impaired performance coupled with slow/non-responding following PC inactivation may highlight the inability to execute the proper actions toward the desired trajectory or goal location.

The present results are also consistent with the notion that the ALT has a role in allocentric spatial encoding (Aggleton & Nelson, 2015; Clark & Harvey, 2016; O’Mara, 2013; Van Der Werf, Jolles, Witter, & Uylings, 2003). There are multiple subdivisions within the anterior thalamic nuclei based upon differences in function and connectivity. For instance, the anteromedial and anteroventral nuclei are thought to be a part of system that synchronizes with hippocampal theta rhythm activity while the anterodorsal nucleus contains a large concentration of HD cells (Taube, 2007; Vertes, Linley, Groenewegen, & Witter, 2015). HD cells are often linked to allocentric spatial processing (Dudchenko et al., 2019; Taube, 1995, 2007) which is supported by observations that experimental manipulation of this neural signal and damage to the ATL produces impairments very similar to hippocampal inactivation or lesions (Aggleton, Hunt, Nagle, & Neave, 1996; Butler et al., 2017). Although the regions that we targeted also included the lateral thalamus in addition to the anterior thalamus, the adjacent laterodorsal thalamus contains HD information while other regions of the lateral thalamic aggregate (centrolateral, lateral mediodorsal nuciel) have also been linked to spatial navigation and memory (Clark & Harvey, 2016; Lopez et al., 2009; Mitchell & Dalrymple-Alford, 2006; Mizumori & Williams, 1993; Perry & Mitchell, 2019; Taube, 1995; Taube & Bassett, 2003).

Though subjectively, it appeared that network inactivation produced changes intermediate to ALT or PC inactivation, in terms of significant differences, the pattern of significant effects was the same for ALT and network inactivations. Thus, the aspects of network inactivation that is consistent with PC inactivation and ALT inactivation (impaired performance) provides further support for the hypothesis that the ALT-PC circuit is critical for transforming spatially relevant contextual demands into the appropriate actions. This transformation would be critical for generating a route to a goal location and executing the proper movements toward the goal (McNaughton, Knierim, & Wilson, 1995; Sutherland & Hamilton, 2004; Wilber et al., 2014). The connections between ALT and PC are largely ipsilateral, so the effect we observed from disconnecting the circuit with contralateral infusions (right PC and left ALT) is consistent with effects observed from circuit disconnection in other regions with similar structural connectivity (Fresno et al., 2019; Jo & Lee, 2010; Wilber et al., 2015).

The place-action task used in this study requires a combination of both allocentric location and egocentric action in order to reach one of four fixed goal locations dependent on the place on the maze. Other “cross maze” task variations (similar maze layout but different task rules) force the animal to utilize a specific strategy (allocentric or egocentric; Aggleton et al., 1996). For these cross-maze variations ALT inactivations produced deficits only when an allocentric strategy was employed, but not when an egocentric strategy was employed (Aggleton et al., 1996). The present task does not distinguish between allocentric heading and allocentric location therefore, animals may be solving the task by transforming a place into action or by transforming a heading into action. Nevertheless, since all inactivation groups produced a similar deficit in the place-action task, these results suggest that the ALT-PC circuit is critical for translating, or at the very least, coordinating between allocentric goal location (or heading) and egocentric action. Future research could further our understanding of allocentric-egocentric coordination by using a paradigm in which there are distinct allocentric, egocentric and transformation components in which allocentric location is dissociated from allocentric heading.

Although the present study found a similar drop in correct responding with PC and ALT inactivations, one notable difference is that following ALT inactivation animals proceeded more quickly to the incorrect location. This could mean that allocentric information was not being translated properly in the absence of the HD signal from ALT, leading to the intact network components generating the incorrect action. The ALT-PC circuit is likely a component of a larger network for coordination of spatial information that includes the RSC and hippocampus (which would both be intact following ALT inactivation). It is important to note that several other regions contribute to egocentric and allocentric spatial information processing. For instance, the hippocampus and entorhinal cortex have specific cell types, place cells and grid cells respectively, that are thought to be the neural substrate of an allocentric cognitive map-like representation of the environment for navigation (Moser, Moser, & McNaughton, 2017; O’Keefe & Nadel, 1978).

The hippocampus has direct connectivity to the RSC, allowing the transfer of hippocampal place information to a brain region that contains a mixture of allocentric and egocentric encoding cells (Wyss & Van Groen, 1992). The RSC also has highly dynamic and adaptive cellular activity that corresponds to a wide range of spatially relevant computations (Alexander & Nitz, 2017). Finally, the RSC sends and receives many projections to both ALT and PC, making it a valuable player in processing egocentric and allocentric spatial information. This is consistent with the hypothesis that HD information from ALT is critical for performing a transformation from allocentric place to egocentric action and that in the absence of this upstream information the transformation of place to action still occurs, possibly by a HPC-RSC-PC circuit, but in the absence of the correct HD leads to incorrect action. Thus it is likely that multiple regions support coordination between ego- and allo-centric representations for navigation. Such network coordination would be essential in order to provide flexibility and efficiency when travelling toward a goal location including the place-action task.

Together the evidence here suggests that the ALT-PC circuit is critical for the coordination between allocentric location and egocentric action in order to reach a goal. Thus the ALT-PC circuit may be critical for *transformation* of allocentric place into egocentric action.

## Acknowledgments

This research was supported by grants from the National Institutes of Health (R01 AG070094 to BJC, and R00 AG049090 and R01 AA029700 to AAW) and Florida Department of Health (FL DOH 20A09).

## Notes

### Competing Interest Statement

The authors have declared no competing interest.

